# Cytosplore-Transcriptomics: a scalable interactive framework for single-cell RNA sequencing data analysis

**DOI:** 10.1101/2020.12.11.421883

**Authors:** Tamim Abdelaal, Jeroen Eggermont, Thomas Höllt, Ahmed Mahfouz, Marcel J.T. Reinders, Boudewijn P.F. Lelieveldt

## Abstract

The ever-increasing number of analyzed cells in Single-cell RNA sequencing (scRNA-seq) experiments imposes several challenges on the data analysis. Current analysis methods lack scalability to large datasets hampering interactive visual exploration of the data. We present Cytosplore-Transcriptomics, a framework to analyze scRNA-seq data, including data preprocessing, visualization and downstream analysis. At its core, it uses a hierarchical, manifold preserving representation of the data that allows the inspection and annotation of scRNA-seq data at different levels of detail. Consequently, Cytosplore-Transcriptomics provides interactive analysis of the data using low-dimensional visualizations that scales to millions of cells.

**Availability:** Cytosplore-Transcriptomics can be freely downloaded from transcriptomics.cytosplore.org

## 1 Introduction

Single-cell RNA sequencing (scRNA-seq) is a valuable technology to identify the cellular composition of complex tissues (Cao *et al.*, 2019). Technological advances over the last decade resulted in a large increase in the acquired data size, scaling to millions of cells, raising major challenges for data analysis (Angerer *et al.*, 2017; Svensson *et al.*, 2018; Lähnemann *et al.*, 2020). Current available tools, such as Seurat (Stuart *et al.*, 2019) and Scanpy (Wolf *et al.*, 2018), provide automated pipelines to analyze scRNA-seq datasets. Although these automated pipelines increase the reproducibility of analyses, they lack the possibility to interactively probe the data and intermediate results, which is essential since often the data analysis is largely exploratory.

Other tools offer interactive visualization and analysis for scRNA-seq data, including ASAP (Gardeux *et al.*, 2017), cellxgene (https://github.com/chanzuckerberg/cellxgene), Granatum (Zhu *et al.*, 2017), Single Cell Explorer (Feng *et al.*, 2019) and UCSC Cell Browser (Speir *et al.*, 2020). However, these tools do not scale to large datasets consisting of millions of cells. In addition, some tools are limited to a list of pre-loaded datasets, and do not allow users to explore, analyze and manually adjust annotations of their own data.

We present Cytosplore-Transcriptomics, a framework for interactive visual analysis and exploration of large scRNA-seq datasets consisting of millions of cells. Building on the principles of Cytosplore (Höllt *et al.*, 2016), we produce a hierarchical representation of the data using HSNE (Pezzotti *et al.*, 2016; Van Unen *et al.*, 2017), which preserves the high-dimensional data manifold. We provide an interactive multiscale exploration of this hierarchy, starting from an abstract embedding containing fewer but representative cells for the global cellular composition, moving to more detailed embeddings of selections of cells on demand. The two-dimensional embeddings of the HSNE hierarchy can be used to cluster and define cell populations at different levels of the hierarchy, or to visualize the expression of selected genes and metadata across cells. Moreover, Cytosplore-Transcriptomics allows an interactive differential gene expression test between selected cell clusters.

## 2 Methods

Cytosplore-Transcriptomics is able to perform data preprocessing, interactive data visualization, as well as downstream analysis such as clustering, cell type annotation and detecting differentially expressed genes across cell groups.

### 2.1 Data input and feature selection

The user can provide data in various formats: (i) csv file containing genes as rows and cells as columns, (ii) hdf5 file including or excluding meta data, (iii) 10X sparse matrix format, or (iv) H5AD file containing a preprocessed Scanpy object. Additionally, meta-data can be uploaded separately in csv format. While uploading the data, CPM (count per million) normalization can be applied, as well as a log(x+1) or square root (sqrt) transformation.

Informative features/genes can be interactively selected to be used for the low-dimensional embedding (Fig. 1A). First, the user may upload a list of genes to exclude from the analysis, such as mitochondrial genes. Next, highly variable genes can be selected by changing the selection threshold applied to the variance. In case of visualizing a previous analysis, it is also possible to upload a list of selected genes to be directly used for the embedding.

**Fig. 1.**
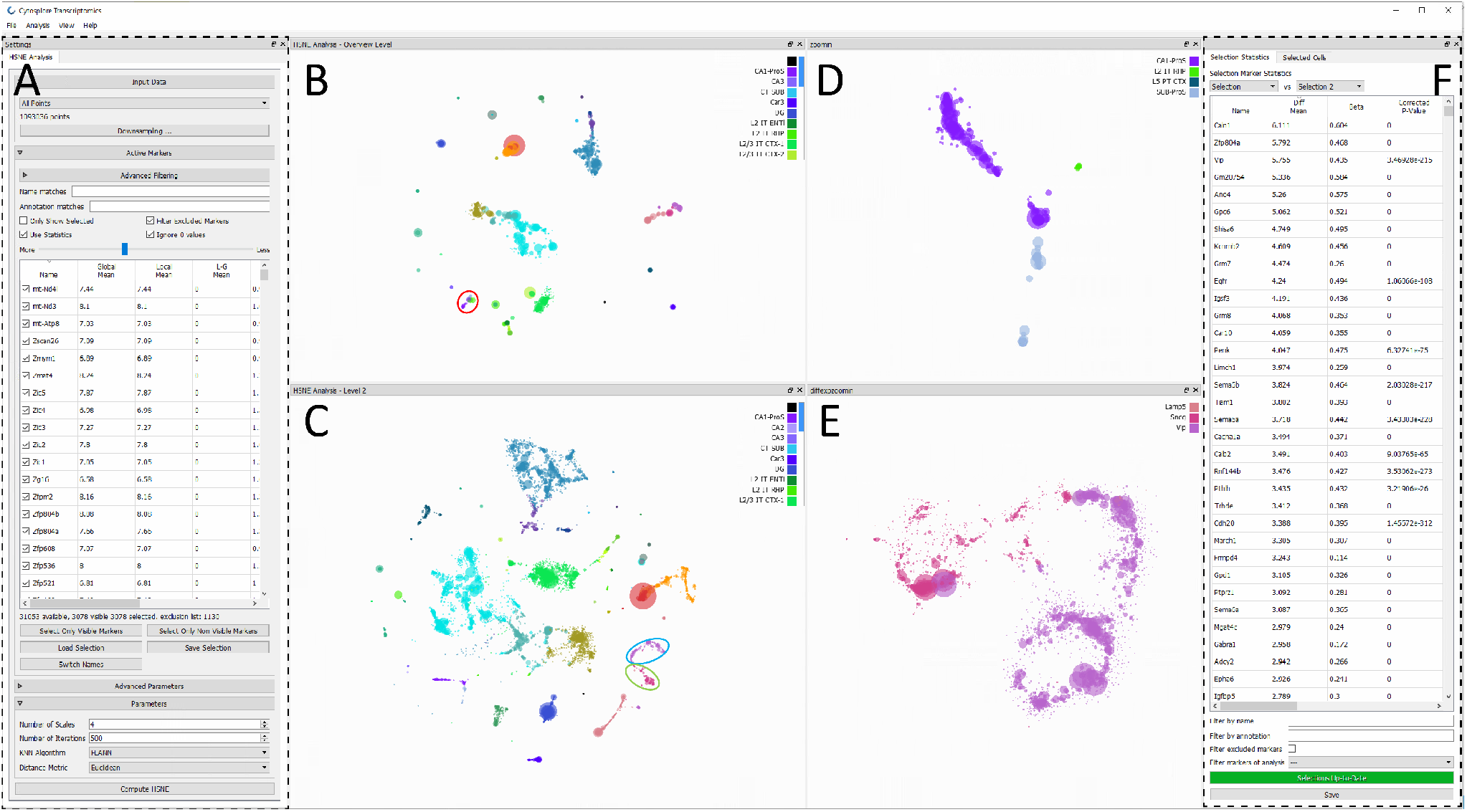
Cytosplore-Transcriptomics software. **(A)** HSNE analysis settings panel, where feature selection can be performed, and HSNE parameters can be selected. **(B)** Exploration of the data hierarchy by first showing the HSNE embedding of the overview scale with only 1,970 cells (0.18% of the total number of cells). **(C)** Zooming one scale deeper into the hierarchy to scale 2 having 11,417 cells (1.04% of the total number of cells). **(D)** HSNE embedding zooming into a specific group of hippocampus cells, highlighted in red in (B), further revealing the cellular diversity within this group. **(E)** HSNE embedding zooming on the Vip and Sncg neurons, used for differential expression analysis, highlighted in blue and green, respectively, in (C). All plots are colored according to the labels from the metadata. **(F)** Differential expression panel showing all genes with their corresponding statistics.

### 2.2 Hierarchical visualization

Once feature selection is performed, a hierarchical low-dimensional embedding of the data can be produced using HSNE. HSNE builds a hierarchy representing the dataset neighborhood in the high-dimensional feature space that preserves the manifold structure of the data, starting from the raw data points moving to multiple abstraction scales in a hierarchical way. The visualization of this hierarchy works in reverse order, by first showing a two-dimensional embedding of the highest scale in the hierarchy (overview scale) containing fewer, but representative, cells. Next, a more detailed embedding can be explored for a selected set of cells, by moving down through the hierarchy. In such a way, HSNE is scalable to millions of cells, without the need of downsampling, with the continuous possibility to explore the data hierarchy at more detailed scales. The number of scales is defined by the user and it is relative to the dataset size, it is recommended to set the number of scales to log10(N/100) where N is the total number of cells in the dataset. At any scale, gene expression and metadata can be overlaid on the low-dimensional embedding.

### 2.3 Clustering and annotation

To define different cell populations in the data, Cytosplore-Transcriptomics provides two different clustering methods, density-based and graph-based clustering. The density-based clustering relies on the low-dimensional embedding, where the layout of the cells indicates the similarity in the high-dimensional feature space. Based on the density representation of the embedding, unsupervised Gaussian Mean Shift (GMS) clustering can be applied to define different cell clusters. On the other hand, graph-based SCHNEL clustering (Abdelaal *et al.*, 2020) can be applied independently from the low-dimensional embedding, as the SCHNEL clustering applies the (Louvain or Leiden) community detection algorithm (Blondel *et al.*, 2008; Traag *et al.*, 2019) on the neighborhood graph of each scale in the hierarchy. Moreover, Cytosplore-Transcriptomics allows the user to manually select and annotate (or correct annotations of) a set of cells of interest.

### 2.4 Differential gene expression

Cytosplore-Transcriptomics provides an interactive differential expression test between different groups of cells (Wilcoxon rank-sum test with Bonferroni multiple testing correction). These groups can be cell clusters or a set of manually selected cells. The set of differentially expressed genes (DEgenes) can be provided either between two groups of cells, or one group versus the remaining cells. Next, the user may pick any of the DEgenes to visualize its expression level on the current embedding.

## 3 Case study

To illustrate the features of Cytosplore-Transcriptomics, we chose the mouse whole cortex and hippocampus dataset from the Allen Institute (https://portal.brain-map.org/atlases-and-data/rnaseq/mouse-whole-cortex-and-hippocampus-10x), representing a relatively large scRNA-seq dataset with over a million cells having diverse cellular populations. We downloaded the original data files, and converted it to one hdf5 file including the metadata (https://doi.org/10.5281/zenodo.4317397). We used Cytosplore-Transcriptomics for visual exploration of this data. First, the data with corresponding metadata is loaded, and we applied CPM normalization and a log(x+1) transformation to the data. Next, we excluded mitochondrial and sex related genes, and selected the top 3,078 highly variable genes for further analysis. An HSNE hierarchy with four scales (data scale + 3 higher scales) was computed and an overview embedding (scale 3) showing only 1,970 cells (0.18% of the full dataset) was visualized (Fig. 1B). This scale shows the overall structure of the dataset with a clear separation between 34 cell populations identified in the metadata. To reveal more detailed structures (Fig. 1C), we zoomed one scale deeper into the hierarchy, examining the embedding of scale 2 comprising 11,417 cells (1.04% of the full dataset). An interesting feature of the hierarchical exploration is the ability to zoom into a specific set of cells. For instance, in Fig. 1D, we focused on a small group of cells from the hippocampus (highlighted in red in Fig. 1B) and generated a separate, more detailed embedding of these cells. This new embedding clearly reveals the heterogeneity in the cellular composition of this specific group of cells, as several smaller subpopulations can be identified from different hippocampal regions, including CA1, retrohippocampal and prosubiculum. Next, we applied the SCHNEL clustering to the 1,970 cells at scale 3, producing 20 cell clusters (Supplementary Fig. S1A). We quantified the agreement of this clustering result with the 34 labels from the metadata using the adjusted Rand index (ARI), measuring the similarity between two different groupings of cells, and obtained an ARI of 0.75 (1 being perfectly similar). We found 10 more clusters when applying SCHNEL to the more detailed scale 2 (Supplementary Fig. S1A), with a total of 30 cell clusters that collectively have an ARI of 0.72 compared to the metadata labels. Finally, we calculated the DE-genes between two adjacent cell populations, Vip and Sncg neurons (highlighted in blue and green, respectively, in Fig. 1C), to reveal the driving genes for these cellular populations. We zoomed into these two populations generating a separate embedding (Fig. 1E), and overlaid the expression of the top DEgenes for each population, showing that *Caln1* is differentially expressed in the Vip neurons, while *Cck* is differentially expressed in the Sncg neurons (Supplementary Fig. S1B). In total, 2,845 DEgenes are obtained (corrected p-value < 0.05, absolute average log2 fold-change > 1), each with their relevant statistics, including mean expression in each population, mean difference between populations, original and corrected p-values (Fig. 1F).

## 4 Conclusions

We developed Cytosplore-Transcriptomics, a standalone tool that facilitates interactive visual exploration and analysis of large scRNA-seq datasets consisting of millions of cells, while preserving the manifold structure of the full data. In addition, it offers many interactive features including, feature selection, clustering and differential gene expression.

