## Supplementary Fig for "Cytosplore-Transcriptomics: a scalable interactive framework for single-cell RNA sequencing data analysis"

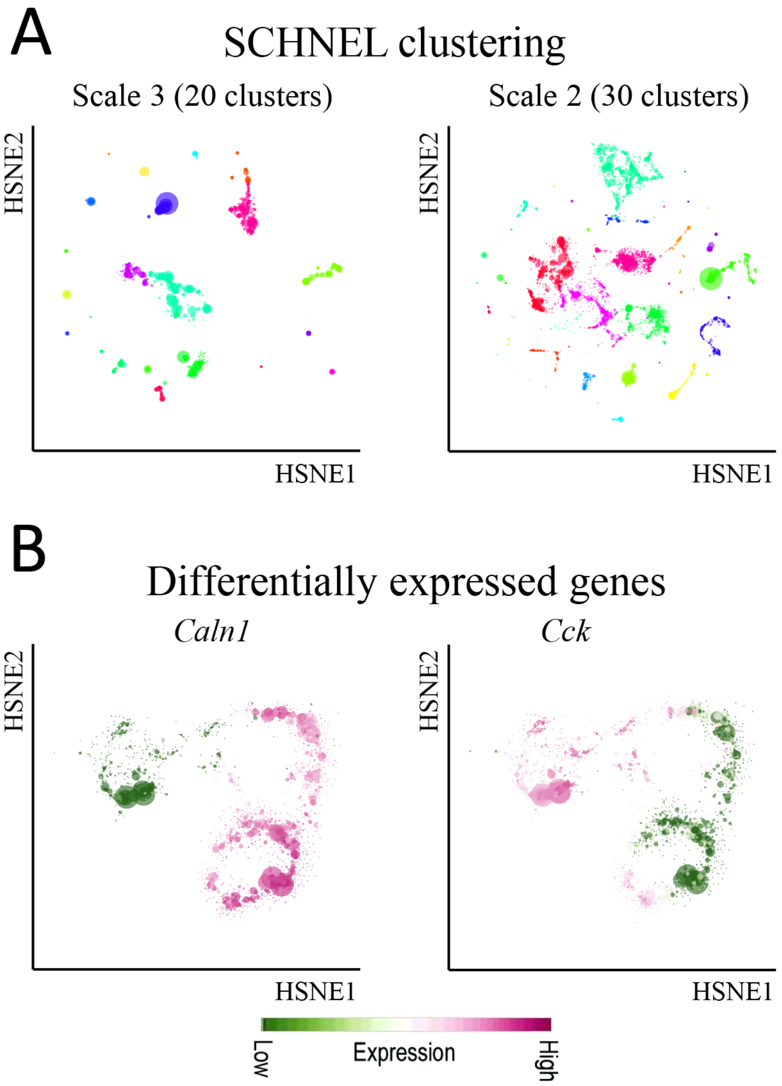


**Supplementary Fig. S1 (A)** SCHNEL clustering of scale 3 (left plot) and scale 2 (right plot) producing 20 and 30 cell clusters, respectively. Colors represent different cell clusters. **(B)** Expression profiles of top DEgenes between two adjacent cell populations, Vip and Sncg neurons, highlighted in blue and green, respectively, in Fig. 1C. The gene expression is overlaid on the HSNE embedding zooming on the two populations of interest (Fig. 1E). *Caln1* is differentially expressed in the Vip neurons, while *Cck* is differentially expressed in the Sncg neurons.
